# Fungi.guru: comparative genomic and transcriptomic database for the Fungi kingdom

**DOI:** 10.1101/2020.06.26.174581

**Authors:** Jolyn Jia Jia Lim, Jace Koh, Jia Rong Moo, Erielle Marie Fajardo Villanueva, Dhira Anindya Putri, Yuen Shan Lim, Wei Song Seetoh, Sriya Mulupuri, Janice Wan Zhen Ng, Nhi Le Uyen Nguyen, Rinta Reji, Margaret Xuan Zhao, Tong Ling Chan, Edbert Edric Rodrigues, Ryan Kairon, Natasha Cassandra Chee, Ann Don Low, Zoe Chen Hui Xin, Shan Chun Lim, Vanessa Lunardi, Fong Tuck Choy, Cherlyn Xin’Er Chua, Kenny Koh Ting Sween, Jonathan Wei Xiong Ng, Marek Mutwil

## Abstract

The fungi kingdom is composed of eukaryotic heterotrophs, which are responsible for balancing the ecosystem and play a major role as decomposers. They also produce a vast diversity of secondary metabolites, which have antibiotic or pharmacological properties. However, our lack of knowledge of gene function in fungi precludes us from tailoring them to our needs and tapping into their metabolic diversity. To remedy this, we gathered genomic and gene expression data of 19 most widely-researched fungi to build a database, fungi.guru, which contains tools for cross-species identification of conserved pathways, functional gene modules, and gene families. We exemplify how our database can elucidate the molecular function, biological process and cellular component of genes involved in various biological processes, by identifying a secondary metabolite pathway producing gliotoxin in *Aspergillus fumigatus*, the catabolic pathway of cellulose in *Coprinopsis cinerea* and the conserved DNA replication pathway in *Fusarium graminearum* and *Pyricularia oryzae*. The database is available at www.fungi.guru.

## INTRODUCTION

The fungi kingdom comprises an enormous diversity of taxa with varied life cycle strategies, ecologies, and morphologies ranging from unicellular yeasts to large mushrooms. Since ancient times, fungi were used as fermenters and food, but recent mycological research demonstrates yet more novel applications of fungi in food production (1, 2). While fungi can cause diseases and decay of building materials and fibers (3), fungi have immensely contributed to agriculture and medicine. For example, secondary metabolites naturally produced by Penicillium mould are vital antibiotics such as penicillin (4), while in agriculture, fungi could become a sustainable alternative for animal feed and revolutionize food processing (5). The sheer diversity of fungal species, estimated at 2.2 to 3.8 million (6), holds much promise for future research.

As more fungi are being discovered and studied, it is becoming clear how little we know about the gene functions in this kingdom (7). Genome sequence alone often cannot reveal the function of the species-specific genes, or how the genes work together to form a biological pathway (8). Genes with similar expression profiles across environmental perturbations, developmental stages, and organs tend to be functionally related, and the identification of these co-expressed genes is thus a powerful tool to study gene function (9–14). These co-expressed genes can be represented as networks, where nodes (or vertices) correspond to genes and edges (or links) connect and indicate genes that have similar expression profiles (15, 16). The analysis of gene expression data and coexpression networks can reveal groups of functionally related genes (i.e., gene modules), while conservation of these networks over large phylogenetic distances can reveal core components of biological processes (10, 17–20). Thus, coexpression analysis is an invaluable tool to reveal gene function (21), but this resource is missing for fungi.

To facilitate the elucidation of gene functions in fungi, we present fungi.guru. This online database allows analysis of coexpression networks, gene expression profiles, ontology terms, gene families, and conserved coexpression clusters of 19 fungal species. The database provides a plethora of tools that allow functional inferences of uncharacterized fungal genes. Our database opens up new possibilities to study fungi and to harness their rich metabolomic arsenal to generate novel drugs and other high-value compounds.

## MATERIALS AND METHODS

### Used genomic and transcriptomic data

Publicly available RNA sequencing experiments for the 20 unicellular and multicellular fungi were identified through NCBI SRA (22). The SRA runtables obtained included experiments from different organs (for multicellular fungi), gene knockout variants, and samples subjected to various chemical and environmental treatments. Coding sequence (CDS) files for each fungus were downloaded from Ensembl Fungi (Table S1) and used to build Kallisto index files for subsequent gene expression estimation with Kallisto v0.46.1 (23). RNA-sequencing experiments for all the fungal species were streamed as fastq files from the European Nucleotide Archive (ENA) (24) by using the LSTRaP-Cloud pipeline (ref). In total, 40843 samples comprising 40.8 terabytes were downloaded for 20 fungi (Table S2). TPM (transcripts per million) gene expression values were generated from fastq files by running the Kallisto quant command in the LSTRaP-cloud pipeline with default parameters (25). For paired-end reads, only the files containing the first read designated with “_1” were downloaded. Single-end and paired-end libraries were mapped with an estimated fragment length of 200 bp and an estimated standard deviation of 20.

To annotate the RNA-seq samples, we used information provided within the SRA runtable and annotation of the sequencing experiments. Annotations include the type of medium the samples were cultivated in, organs, location, sexual/asexual types, temperature, and strains. The annotations were used to construct expression profiles on the CoNekT database (26).

### Quality control of RNA-seq experiments and construction of coexpression networks

For each fungal species, RNA-seq samples were quality-controlled by removing samples that showed low absolute number of processed reads (NPR) and low percentage of pseudoaligned reads (PPR) to the CDS (denoted by Kallisto as n_pseudoaligned and p_psuedoaligned respectively)(25, 27). The thresholds were set to remove samples with <1 million NPR and a species-specific threshold of PPR. The PPR threshold was estimated by manual inspection of scatter plots, which show NPR and PPR values on the x- and y-axis, respectively (Figure S1). Fungal species *Ramularia collo-cygni* showed low pseudoalignment values and was eliminated from the project. RNA-sequencing samples that passed the thresholds were used to construct expression matrices for each species. In total, 21135 out of 40843 samples passed the quality control (Table 1, Table S2).

**Table 1.**
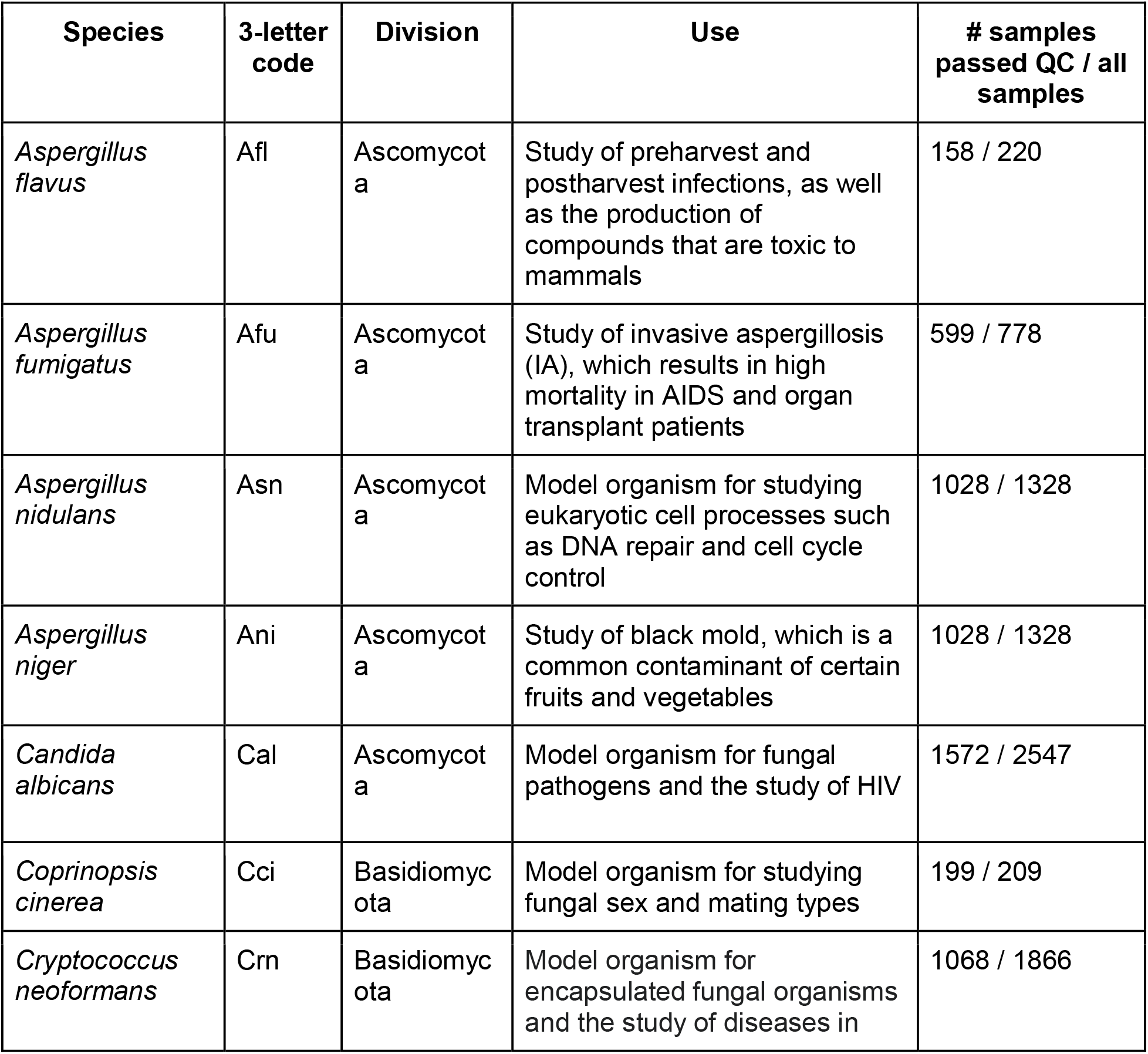
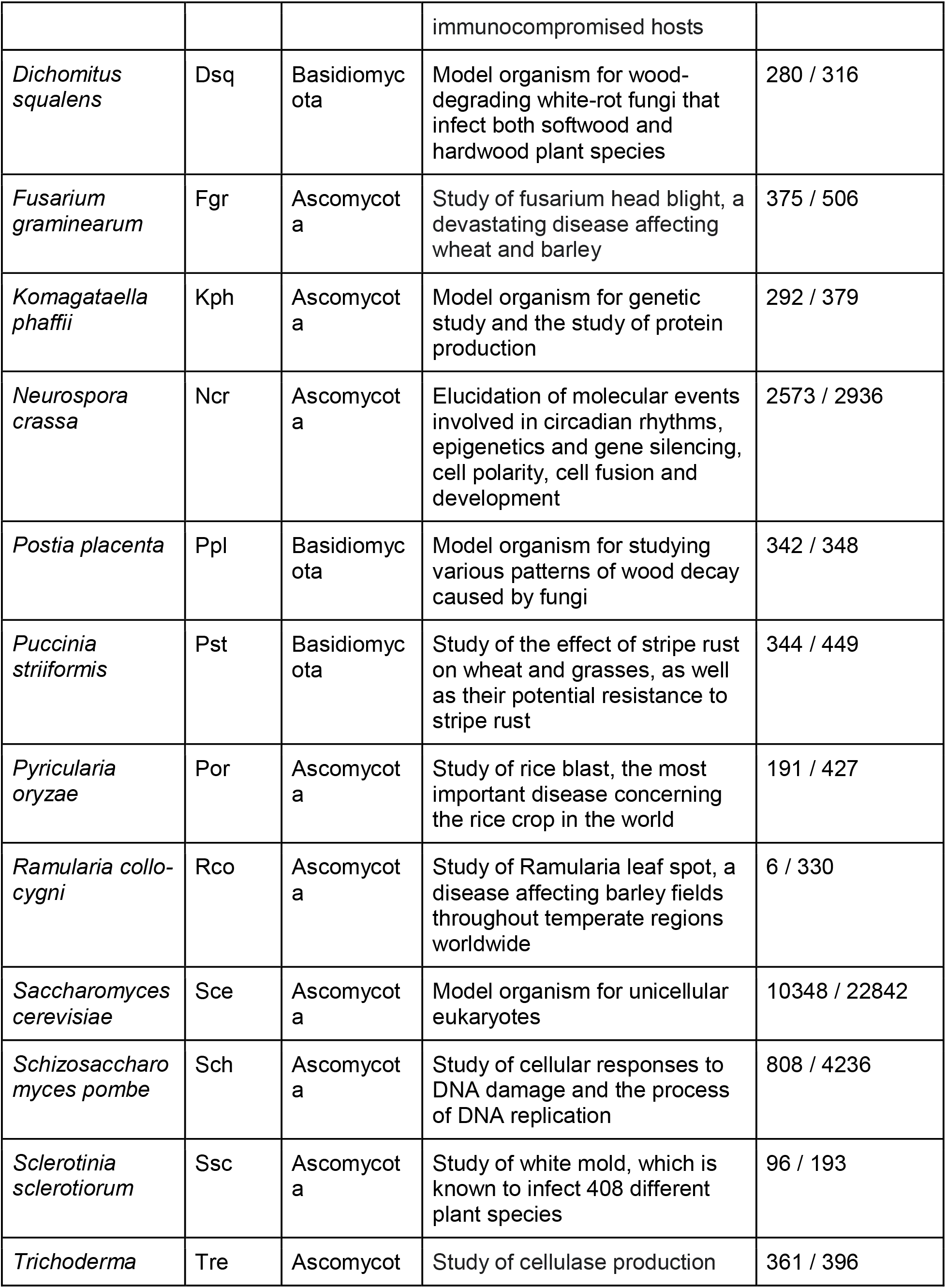
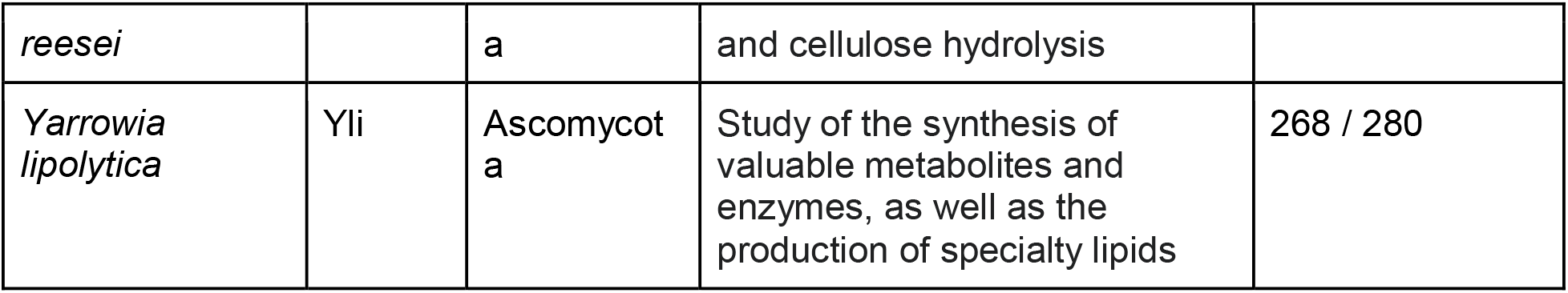
Fungal species included in the database.

### Functional annotation of fungal coding sequences

The protein IDs were obtained from the pep files available on Ensembl Fungi. Interproscan-5.44-79 (28) was used to obtain the Pfam domains and Gene Ontology terms (GO terms) for each protein in the 20 species. The groups of orthologous genes were obtained using Orthofinder v2.3.12 (29) using Diamond (30) with default settings.

### Construction of fungi.guru database

The coexpression networks were constructed by using the Highest Reciprocal Rank metric (31). Conekt database framework with default settings was used to construct the database (26). Coexpression clusters for each species were generated via Heuristic Cluster Chiseling Algorithm (HCCA)(31), where cluster sizes were limited to 100 genes.

## RESULTS

To allow comparison of the coexpression relationships across the fungi kingdom, the genes have been assigned to Pfam domains (via InterproScan)(28) and gene families (via OrthoFinder)(29). Our database offers multiple ways to query it, that can be found in the ‘Search’ panel on the top of the page. The user can find the gene of interest by using gene IDs (e.g., *CBF69394*), sequence similarity (BLAST), Pfam domains, or Gene Ontology terms.

The database allows viewing the genomic and transcriptomic data on multiple levels. For example, it contains pages for species (www.fungi.guru/species), genes (www.fungi.guru/sequence/view/20959), gene families (www.fungi.guru/family/view/117804), coexpression clusters (http://www.fungi.guru/cluster/view/741), neighborhoods (www.fungi.guru/network/graph/15360), phylogenetic trees of families (www.fungi.guru/tree/view/45802), Pfam domains (www.fungi.guru/interpro/view/5689), Gene Ontology terms (www.fungi.guru/go/view/16530) and others. These pages contain data relevant to the type of the viewed data. For example, gene pages contain the functional annotation, the cDNA and protein sequences, assigned gene family, phylogenetic tree of the family, expression profiles, coexpression neighborhood and cluster, significantly similar neighborhoods, and Gene Ontology information. Conversely, Gene Ontology pages indicate the annotation of the GO term, the number of genes in the 19 fungi that have the GO term, and the co-expressed clusters that are enriched for the genes with this GO. A full description of the features is found at localhost/features.

To exemplify these features, we provide three typical case studies addressing different aspects of fungal biology.

### Example 1: Identification of specialized metabolism hubs by coexpression analysis

Coexpression analysis has been used successfully to study numerous metabolic pathways in plants (32, 33) and fungi (refs). To exemplify how our database can be used to study specialized metabolism in fungi, we first set to identify ‘backbone’ enzymes responsible for the biosynthesis of the different metabolite classes. To this end, we used Secondary Metabolite Unique Regions Finder (SMURF) database (34), to assign the genes of the 19 fungi to nonribosomal peptide synthases (NRPSs), polyketide synthases (PKSs), NRPS-like enzymes, PKS-like enzymes, Hybrid enzymes and dimethylallyl tryptophan synthase (DMAT). The analysis revealed that the 19 fungi have a diverse set of the backbone enzymes, ranging from 80 enzymes in *Aspergillus niger* (Ani), to 0 in *Cryptococcus neoformans* (Crn), where five (Asn, Ani, Afu, Por, Afl, Table 1) of the 19 fungi were shown to have more than 40 backbone enzymes (Figure S2A).

To identify groups of genes involved in metabolite synthesis and conditions that result in high metabolite production, backbone enzymes were first retrieved from SMURF (34), and entered into ‘Tools/Create custom network’ tool (www.fungi.guru/custom_network/), which identifies coexpression relationships in the provided list of genes. The custom network analysis of specialized metabolism genes in *Aspergillus fumigatus* produced an interactive network that revealed a high degree of coexpression among the backbone enzymes, indicating that biosynthesis of the various metabolites is transcriptionally coordinated (Figure 1A).

**Figure 1.**
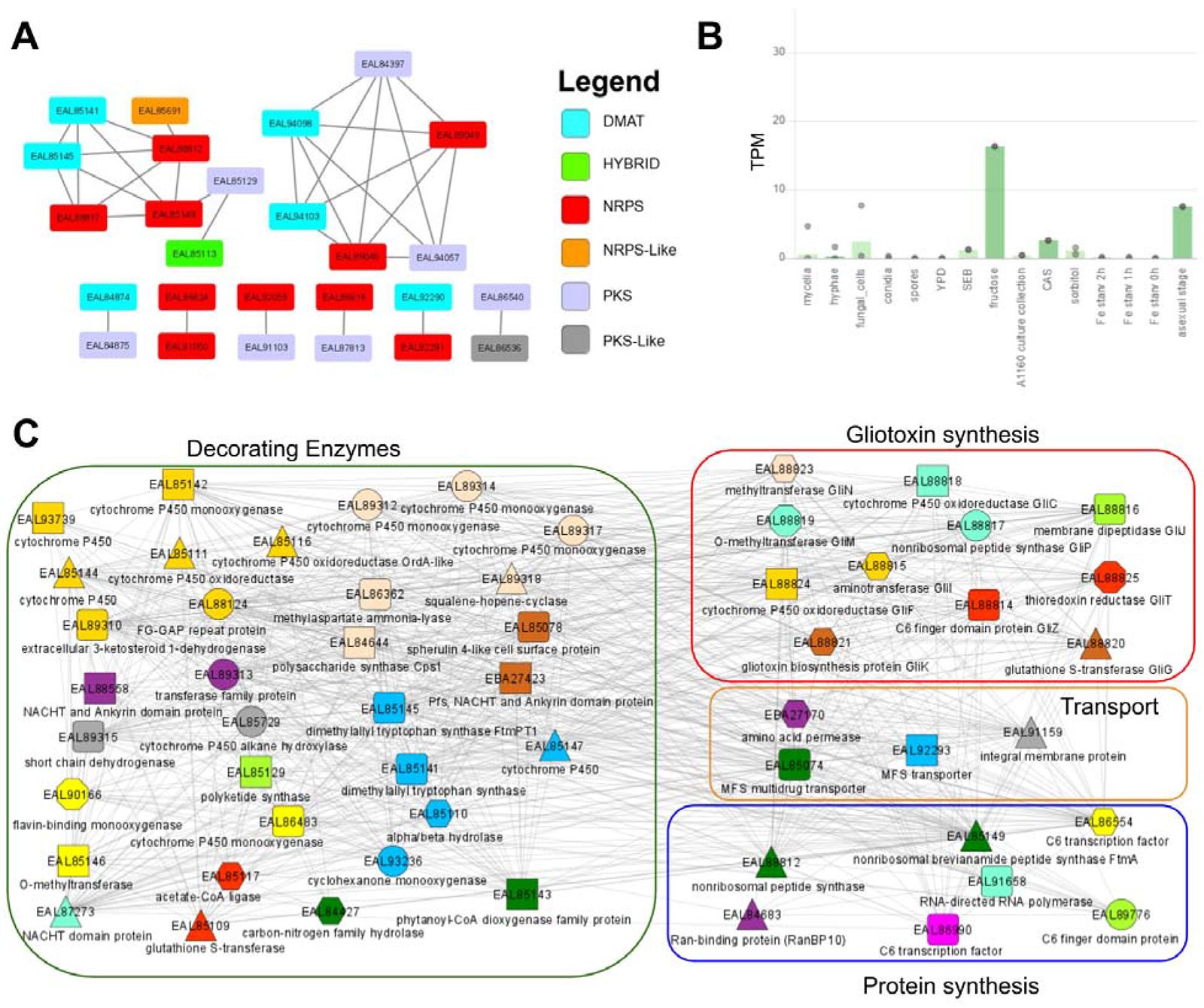
Coexpression analysis of specialized metabolic hubs in *Aspergillus fumigatus*. A) Coexpression network of the backbone enzymes involved in specialized metabolism. Nodes represent genes, while edges connect co-expressed genes. For brevity, enzymes that are not co-expressed are not shown. The color of the node indicates the type of backbone enzyme. The plot was generated by exporting the network file and opening the file in standalone Cytoscape. B) Expression profile of *EAL85149*, a nonribosomal peptide synthase ftmA that shows the highest connection to other backbone enzymes. The different samples are represented on the x-axis, while the gene expression values in the form of Transcripts Per Million (TPM) are indicated on the y-axis. The bars and the dots indicate the average and the minimum/maximum values of the RNA-seq experiments in the sample, respectively. C) Coexpression network of *EAL85149*. Nodes represent genes, while edges connect co-expressed genes. Colored shapes indicate which genes contain Pfam domains and orthogroups in common. For brevity, only part of the network is shown. The network contains four types of genes: gliotoxin family genes (indicated in the red rectangle), genes involved in protein synthesis (blue rectangle), metabolite transporters (orange rectangle), and decorating enzymes (green rectangle). The function of the genes can be inferred from the legend and by clicking on the nodes.

One of the *Aspergillus fumigatus* backbone enzymes, *EAL85149*, nonribosomal peptide synthase *ftmA*, was found to be co-expressed with a large variety of genes involved in secondary metabolite synthesis (Figure 1A), suggesting that the enzyme is part of a transcriptional program that produces numerous metabolites. By entering the EAL85149 into the search box, we arrived at a gene page dedicated to EAL85149 (www.fungi.guru/sequence/view/20959). The expression profile (www.fungi.guru/profile/view/27005) revealed that the gene shows the highest expression in fructose medium (Figure 1B). Since specialized metabolism tends to be transcriptionally regulated (35, 36), this information can be used to identify growth conditions needed to induce high production of specialized metabolites.

*EAL85149* is involved in the biosynthesis of toxins such as gliotoxins and brevianamide F, a precursor bioactive prenylated alkaloid (37). Brevianamide F acts as a precursor for fungal toxins fumitremorgin C and trypacidin that can prevent phagocytosis of fungal conidia, and serves an essential protective role for the pathogenesis of fungi (38). To identify genes that are functionally related to *EAL85149*, we viewed its coexpression network available at the gene page (www.fungi.guru/network/graph/15360)(Figure 1C). 11 out of 13 gliotoxin biosynthesis genes are found in the gliotoxin gli coexpression cluster, with the exception of *GliA*, responsible for the exportation of extracellular gliotoxin essential for gliotoxin tolerance (39) and *GliH* whose function is still unknown (40). Interestingly, the gliotoxin biosynthesis genes found in the coexpression cluster are also part of the genomic gli cluster, where the gli biosynthetic genes are found on the same chromosomal locus (41). The coexpression cluster also contains a variety of transporters such as Major Facilitator Superfamily (MFS) transporter that could be used to secrete gliotoxin and other metabolites potentially synthesized by the coexpression cluster. Additionally, the coexpression cluster contains large numbers of cytochrome P450s, which are decorative enzymes essential for secondary metabolite synthesis (42).

In addition to the discussed genes, we observed multiple genes coding for hypothetical proteins in the cluster. Since these genes are co-expressed with the gli pathway, they could also play a crucial role in secondary metabolite synthesis and merit further study. Thus, this example shows how fungi.guru can be easily used to reveal conditions that are likely to control the production of metabolites and reveal novel genes involved in the metabolite production (Figure S3 shows a similar analysis performed in *Aspergillus flavus*).

### Example 2: Gene Ontology search reveals clusters important for cellobiose degradation

Many fungi, especially saprobic fungi, obtain energy from decaying organic matter. Their main source of energy comes from digesting decomposing plant material such as cellulose and lignin (43). Basidiomycetous fungi, for example, *Coprinopsis cinerea* (*Cci*)(44), are one of the most potent degraders of cellulose as they often grow in environments that are cellulose-abundant, such as dead wood and plants (45, 46). Plant cell walls are multi-layered and made up of different materials which include cellulose, hemicelluloses, and lignins (47), leading to the need for fungi to produce various types of hydrolytic enzymes (endoglucanase, cellobiohydrolase, and beta-glucosidase) to break down these biomolecules into substances that can be used as sources of energy for cellular activity (48). Thus, depending on the composition of the organic matter decomposed, different fungal species utilize their own unique set of enzymes and metabolic pathways for cellobiose degradation (49).

To gain insight into cellobiose degradation in *Cci*, we entered ‘cellulose catabolism’ into ‘Tools/Find enriched clusters’ (www.fungi.guru/search/enriched/clusters), which revealed Gene Ontology term GO:0030245 (GO term for Cellulose Catabolic Process). The database identified six clusters significantly enriched (p-value for enrichment < 0.05) for this GO process in six species (*Aspergillus nidulans*, *Aspergillus niger*, *Coprinopsis cinerea*, *Neurospora crassa*, *Pyricularia oryzae*, and *Sclerotinia sclerotiorum*). For *Cci*, we focused on cluster 21, which was enriched for GO:0030245 (p-value for enrichment < 0.001). Using the database, we arrived at a page that details the cluster analysis of genes involved in cellobiose degradation in *Cci* (www.fungi.guru/cluster/view/741)(Figure 2A).

**Figure 2.**
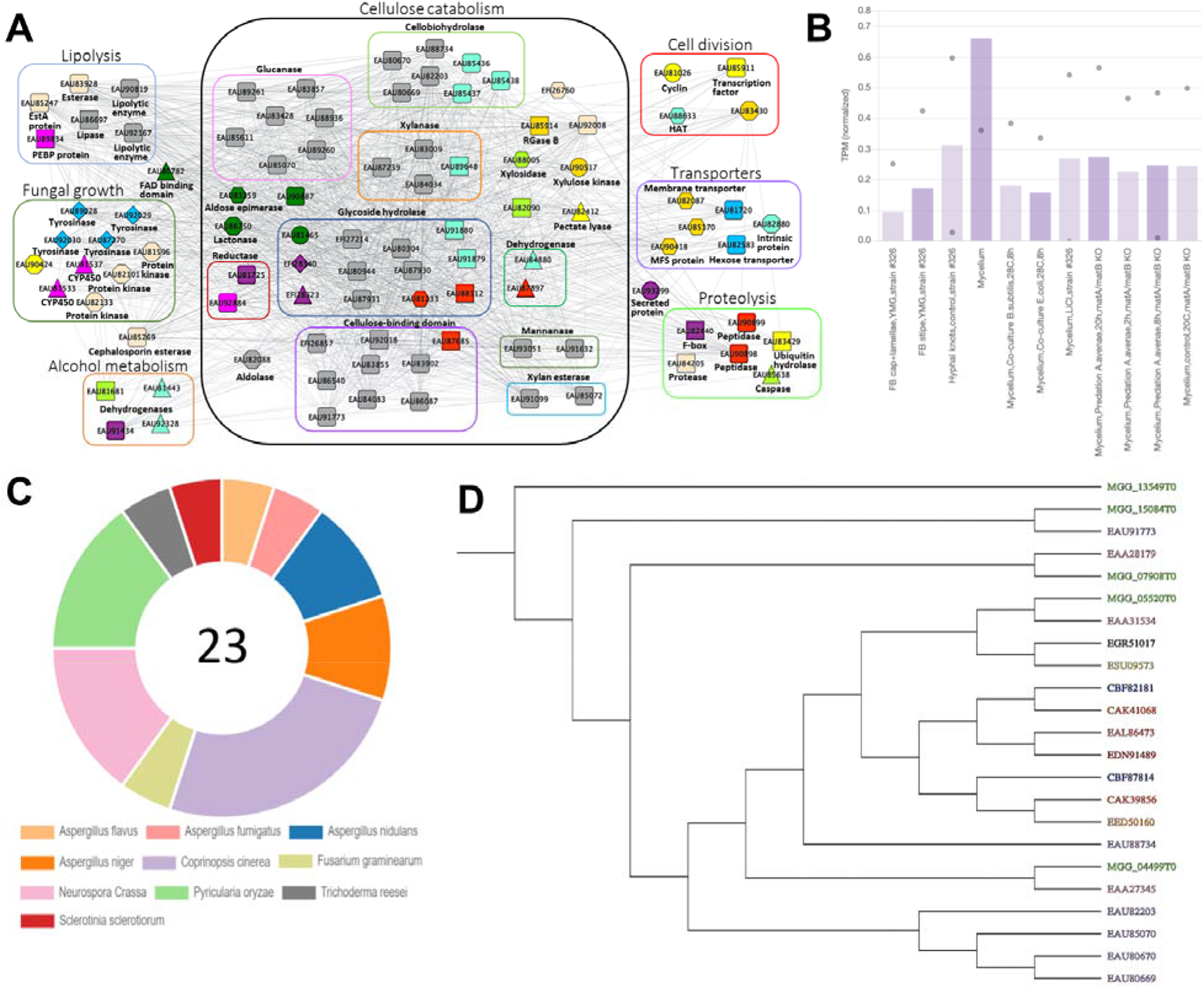
Cluster analysis involved in cellulose catabolic process in *Coprinopsis cinerea*. A) Coexpression network of genes identified as enriched for GO:0030245 (GO term for Cellulose Catabolic Process). Nodes represent genes, while co-expressed genes are connected by edges. For brevity, novel genes (hypothetical proteins) are not shown in the network. Colored shapes indicate which genes contain Pfam domains and orthogroups in common. B) Average expression profile of cluster 21 for *Cci*. Only part of the expression profile containing the discussed sample is shown. The different samples are represented on the x-axis, while the gene expression values in the form of Transcripts Per Million (TPM) are indicated on the y-axis. The bars and the dots indicate the average and minimum/maximum values of the RNA-seq experiments in the sample. C) The proportion of sequences in the cellobiohydrolases Gene Family: OG_03_0001111. There are a total of 23 sequences with this label, from 10 different species. D) Phylogenetic tree of cellobiohydrolases gene family OG_03_0001111.

The coexpression cluster contained 150 genes, which were involved in different processes (alcohol metabolism, cellulose catabolism, cell division, fungal growth, lipolysis, proteolysis, and transporters)(Figure 2A). We observed that cellobiohydrolases (50), cellulose-binding domains (51), glucanases (52), glycoside hydrolases (53), xylanases (54), and sugar transporters are the main genes present in the cluster, supporting the function of this cluster in cellobiose degradation and uptake of sugars. The cluster page also contains information about the enriched GO terms, whether similar clusters can be found in other fungi, presence of InterPro domains, enriched GO terms, and presence of gene families. Additionally, we also investigated the expression profile for this cluster (Figure 2B) and observed that the gene expression is the highest in the mycelium, suggesting that these structures are most active in degrading cellulose in *Cci*.

The most abundant gene family in cluster 21 was OG_03_0001111, which contains genes annotated as cellobiohydrolases. We clicked on OG_03_0001111 to arrive at a page dedicated to the family (www.fungi.guru/family/view/117804). The gene family page revealed that *Cci* contains the highest proportion of cellobiohydrolases (Figure 2C), suggesting that it is one of the more potent degraders of cellobiose and its cellobiohydrolase gene family may have undergone a higher degree of expansion compared to the other species. The page also contains a link to the phylogenetic tree generated by Orthofinder (www.fungi.guru/tree/view/45802). The tree (Figure 2D) shows us that the cellobiohydrolase family has expanded in *Cci* and this could hypothetically be attributed to the further enhancement of cellobiose degradation ability in *Cci*.

Since we identified multiple such clusters in different fungi (please see a similar analysis done in *Postia placenta*, Figure S4), we propose that our database is a good starting point to study cellulose degradation in fungi. Thus, the database allows easy identification of coexpression clusters, genes, and gene families relevant to the biological process of interest.

### Example 3. Comparative analyses of DNA replication in fungi

Coexpression relationships can be conserved across species, where coexpression clusters containing the same gene families and Pfam domains are found in multiple species (10, 11, 55). Identifying these conserved coexpression clusters thus enables the identification of conserved biological pathways and the core genetic components of these pathways (ref). Fungi.guru contains multiple tools that allow the identification of the conserved pathways.

To exemplify these comparative features, we examined the process of DNA replication between 2 fungi known for their infectious capabilities - *Fusarium graminearum* and *Pyricularia oryzae*. Under ‘Tools/Find enriched clusters’, we searched for enriched clusters using the GO term “DNA replication” with *F. graminearum* selected as target (www.fungi.guru/search/enriched/clusters). By clicking “Show cluster”, the database revealed that Cluster_10 in *F. graminearum* is enriched for this term (www.fungi.guru/cluster/view/1203). Under ‘Similar Clusters’ table, we identified *P. oryzae* cluster 10 as being similar to *F. graminearum* cluster 10 with Jaccard index value of 0.202 (26). Clicking on the ‘Compare’ button shows the coexpression networks of these two conserved modules, where only genes that have a homolog or Pfam domain in both clusters are shown (Figure 3A).

**Figure 3.**
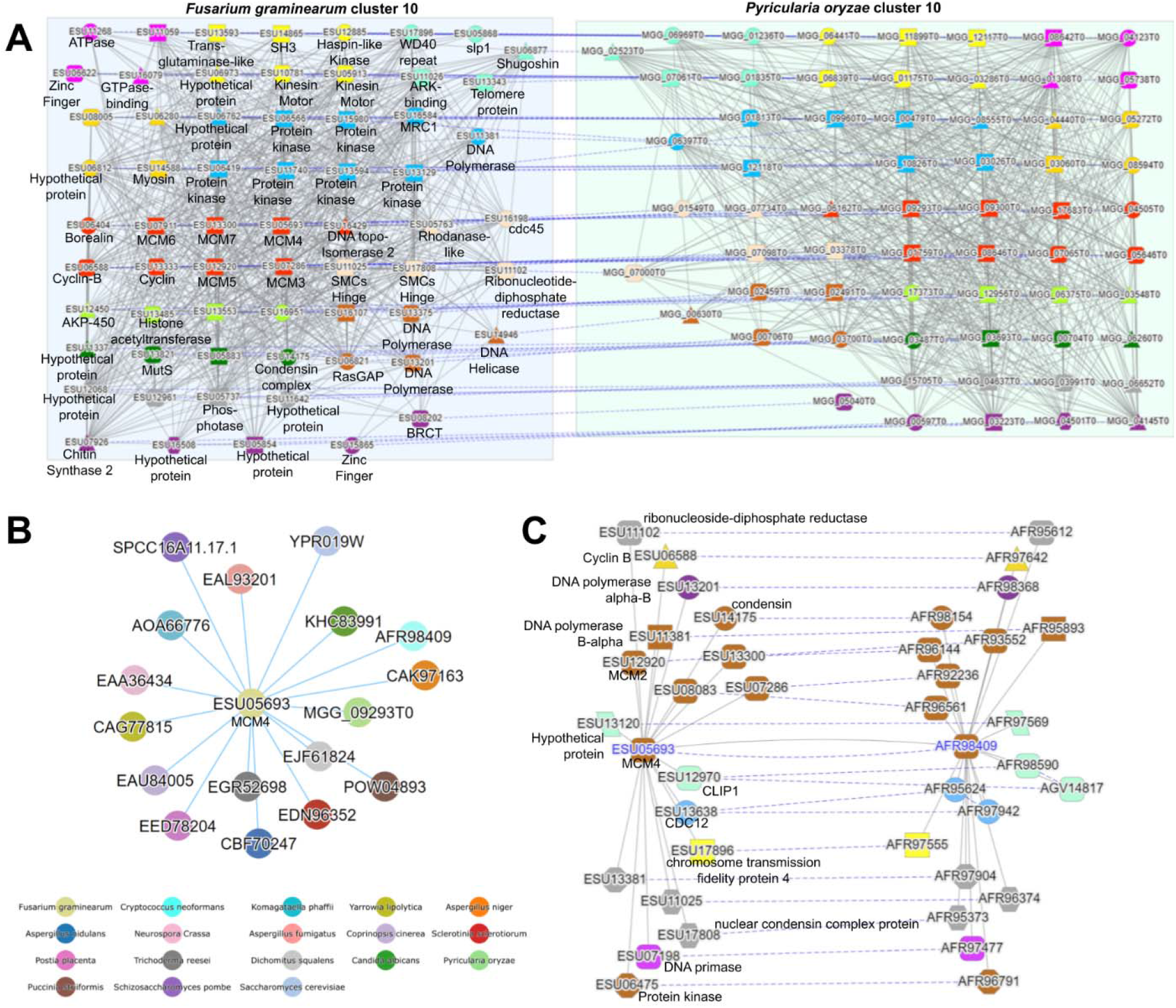
Comparative analysis of gene coexpression networks of DNA replication in *F. graminearum* and *P. oryzae*. A) Comparative analysis of conserved coexpression clusters. Nodes represent genes, while co-expressed genes are connected by edges. The color and shape of the nodes indicates the gene family and pfam domains of the genes. The dashed edges between the conserved clusters connect genes that belong to the same gene families, while gray edges connect co-expressed genes. B) Expression context conservation network of *ESU0569*. Each node indicates a gene co-expression neighborhood, while blue edges connect conserved neighborhoods. The color of the nodes indicates the fungal species. C) Conserved co-expression neighborhoods. Please see text in panel 3A for legend.

As expected for clusters significantly enriched for GO term DNA replication, the genes in the conserved clusters are involved in chromatin regulation and DNA replication. The genes belong to multiple conserved gene families, including histone acetyltransferase, minichromosome maintenance protein complex (MCM), DNA polymerase, and others (Figure 3A). Histone acetyltransferases are involved in the loosening of the condensed chromatin structure to allow DNA replication machinery access to the template (56). Cyclin-B activates downstream cyclin-dependent kinases involved in the M phase of the cell cycle, while other cyclin types control kinases for other stages in the cell cycle (57). MCM is known to be involved in the assembly of the replication initiation complex at the origin of replication (58). Cdc45 form the catalytic core of DNA helicase that serves to unwind the double-stranded DNA (59). Type 2 DNA topoisomerase introduce negative supercoils in the DNA to release the stress on the DNA set by the unwinding action of DNA Helicase (60).

In addition to identifying conserved clusters, our database can also reveal which gene coexpression neighborhoods are similar. For example, *ESU05693* (www.fungi.guru/sequence/view/89556) is a DNA replication licensing factor mcm4 that belongs to OG_03_0001507 gene family (www.fungi.guru/family/view/118200). The gene page for *ESU05693* contains a table ‘Expression context conservation’ which lists other genes from OG_03_0001507 gene family that have a coexpression neighborhood that contains similar content of gene families to *ESU05693*. By clicking on ‘View ECC as graph’, the database indicated that 17 genes from the OG_03_0001507 gene family belonging to 17 different species have similar coexpression neighborhoods (Figure 3B). This is not surprising as DNA replication is a well-conserved process. The contents of the co-expressed neighborhoods can be viewed by clicking on the ‘View ECC pair as graph’ in the ‘Expression context conservation’ table. Similarly to the comparative analysis of the clusters (Figure 3A), the database displays the contents of the two neighborhoods and highlights the conserved gene families and Pfam domains (Figure 3C). The contents of the two coexpression neighborhoods are also related to DNA replication. Thus, fungi.guru allows identifying conserved coexpression clusters and neighborhoods across species, which can be invaluable to identify novel genes and pathways relevant to a biological pathway of interest.

## CONCLUSION

To remedy the paucity of functional information for fungi, we constructed www.fungi.guru, a user-friendly database that facilitates the analysis of genomic and transcriptomic data. The tools available in the database allow for the identification and analysis of co-expressed gene neighborhoods, clusters, and gene families. While our examples addressed specialized metabolism, cellulose degradation, and cell division, the database can be used to study molecular function, biological process, and cellular component of all genes expressed in the 19 fungi present in the database. We envision that the gene function prediction tools present in fungi.guru will enhance the selection of relevant genes to further functional analyses.

## DATA AVAILABILITY

The expression matrices, RNA-seq sample annotation and gene families are available from the Supplementary Data. The co-expression networks, coding and protein sequences can be downloaded from www.fungi.guru.

## ACKNOWLEDGEMENTS

We would like to acknowledge Dr. Irene Julca nad Dr. Riccardo Delli-Ponti for their help in setting up OrthoFinder and Interproscan analyses. William Goh is acknowledged for helping in the initial steps of the project. Finally, we would like to thank Ryan Chee Khiang Ng for techical assistance.

## SUPPLEMENTAL TABLES

**Table S1. Genome versions used to build the database**

**Table S2. Pseudoaligning statistics of the 20 fungal species.**

**Table S3. Expression profile of ASPFL genes across RNA-Seq experiments** Rows contain transcripts per million (TPM) expression values for a particular gene, while columns contain all expression values captured by an RNA-Seq experiment. The first row contains all the RNA-Seq experiment IDs while the first column contains all the gene IDs of the fungi.

**Table S4. Expression profile of ASPFU genes across RNA-Seq experiments**

**Table S5. Expression profile of ASPNI genes across RNA-Seq experiments**

**Table S6. Expression profile of ASPNID genes across RNA-Seq experiments**

**Table S7. Expression profile of CANAL genes across RNA-Seq experiments**

**Table S8. Expression profile of COPCI genes across RNA-Seq experiments**

**Table S9. Expression profile of CRYNE genes across RNA-Seq experiments**

**Table S10. Expression profile of DICSQ genes across RNA-Seq experiments**

**Table S11. Expression profile of FUSGR genes across RNA-Seq experiments**

**Table S12. Expression profile of KOMPH genes across RNA-Seq experiments**

**Table S13. Expression profile of NEUCR genes across RNA-Seq experiments**

**Table S14. Expression profile of POSPL genes across RNA-Seq experiments**

**Table S15. Expression profile of PUCST genes across RNA-Seq experiments**

**Table S16. Expression profile of PYROR genes across RNA-Seq experiments**

**Table S17. Expression profile of SACCE genes across RNA-Seq experiments**

**Table S18. Expression profile of SCHPO genes across RNA-Seq experiments**

**Table S19. Expression profile of SCLSC genes across RNA-Seq experiments**

**Table S20. Expression profile of TRIRE genes across RNA-Seq experiments**

**Table S21. Expression profile of YARLI genes across RNA-Seq experiments**

## SUPPLEMENTAL FIGURES

**Figure S1.**
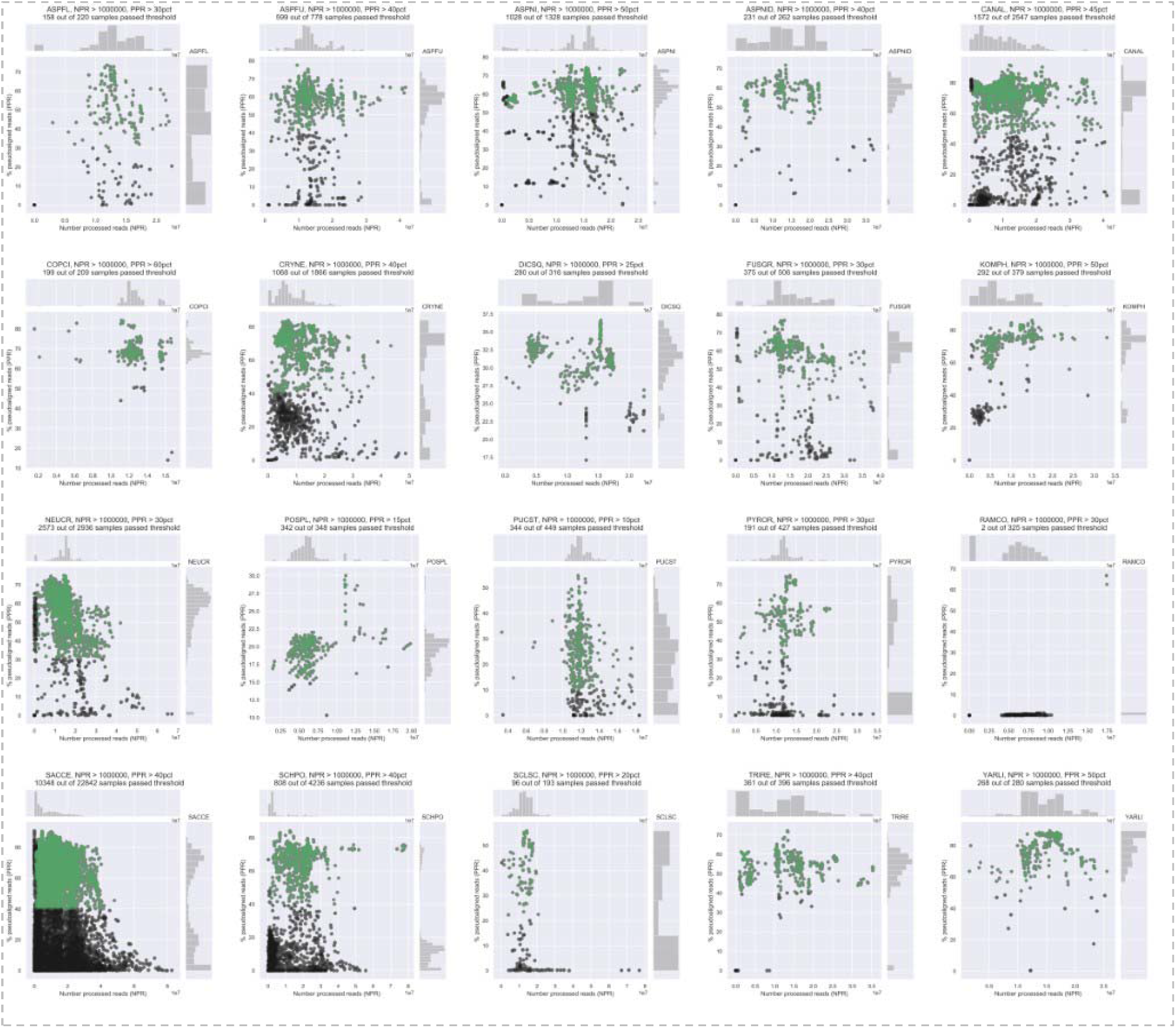
Quality control and data selection for the 20 fungi. The number of processed reads (NPR, x-axis) versus the percentage of pseudoaligned reads (PPR, y-axis) are shown for all 20 species, which are arranged in alphabetical order. Each point represents one sample, where black and green indicate samples that failed and passed the quality control, respectively. The histograms to the right and on top indicate the distributions of the PPR and NPR values, respectively. The sample quality control outcomes are shown in Table S1.

**Figure S2.**
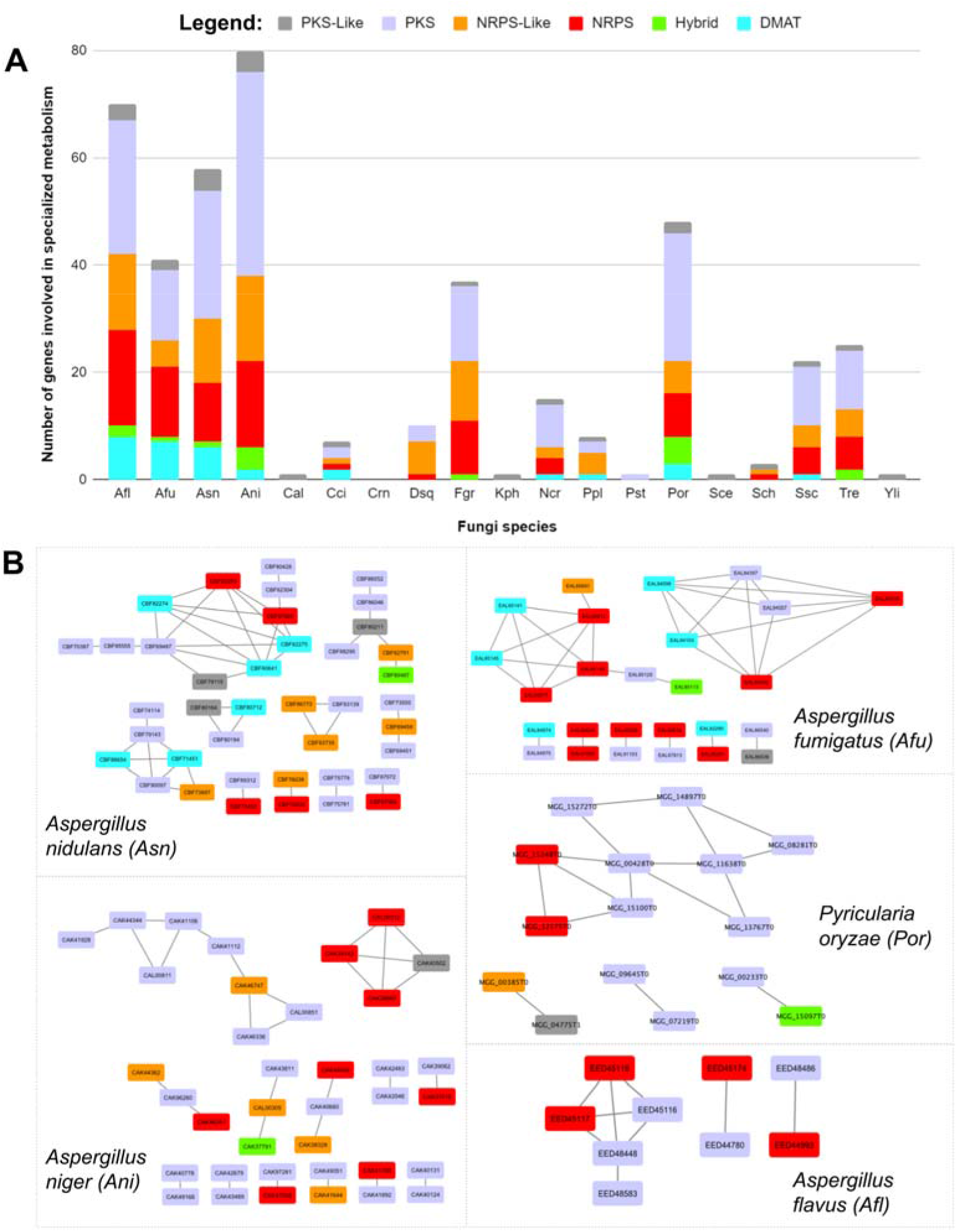
Analysis of specialized metabolism in the 19 fungi. A) The bar chart indicates the fungi (x-axis) and the number of genes (y-axis) predicted to be involved in specialized metabolism by SMURF. The types of metabolites are indicated by color. B) This panel presents five fungal species that have diverse networks of backbone genes as predicted by SMURF.

**Figure S3.**
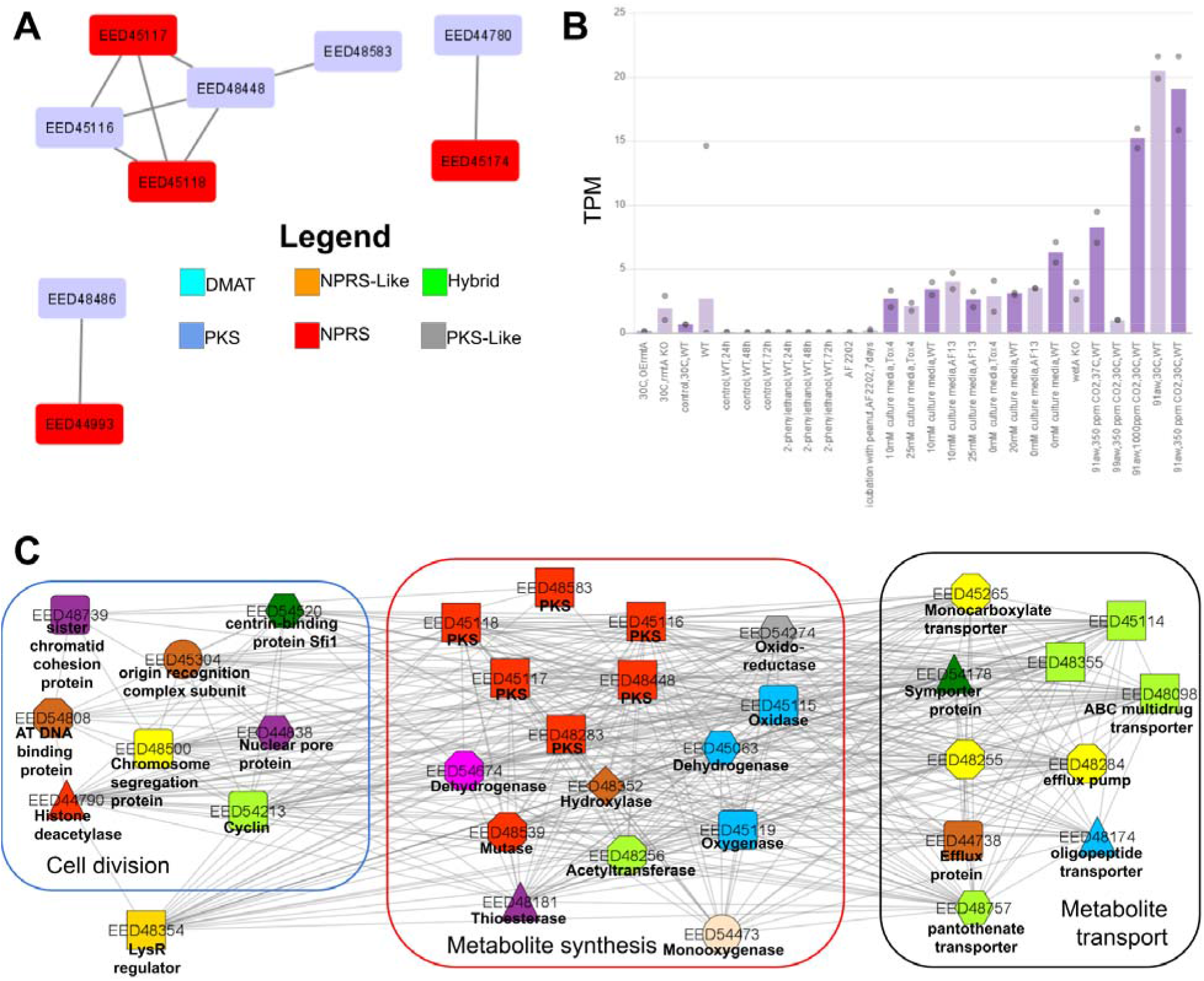
Analysis of specialized metabolic hubs in *Aspergillus flavus*. A) Coexpression network of the backbone enzymes involved in specialized metabolism. Nodes represent genes, while co-expressed genes are connected by edges. For brevity, enzymes that are not coexpressed are not shown. The color of the node indicates the type of backbone enzyme. B) Expression profile of *EED48448*, a putative polyketide synthase that shows the highest connection to other backbone enzymes. The different samples are represented on the x-axis, while the gene expression values in the form of Transcripts Per Million (TPM) are indicated on the y-axis. The bars and the dots indicate the average and the minimum/maximum values of the RNA-seq experiments in the sample, respectively. C) Coexpression network of *EED48448*. Nodes represent genes, while edges connect co-expressed genes. Colored shapes indicate which genes contain Pfam domains and orthogroups in common. For brevity, only part of the network is shown. The network contains three types of genes: metabolic enzymes (indicated in the red rectangle), cell division genes (blue rectangle), and metabolite transporters (black rectangle). The function of the genes can be inferred from the legend and by clicking on the nodes.

**Figure S4.**
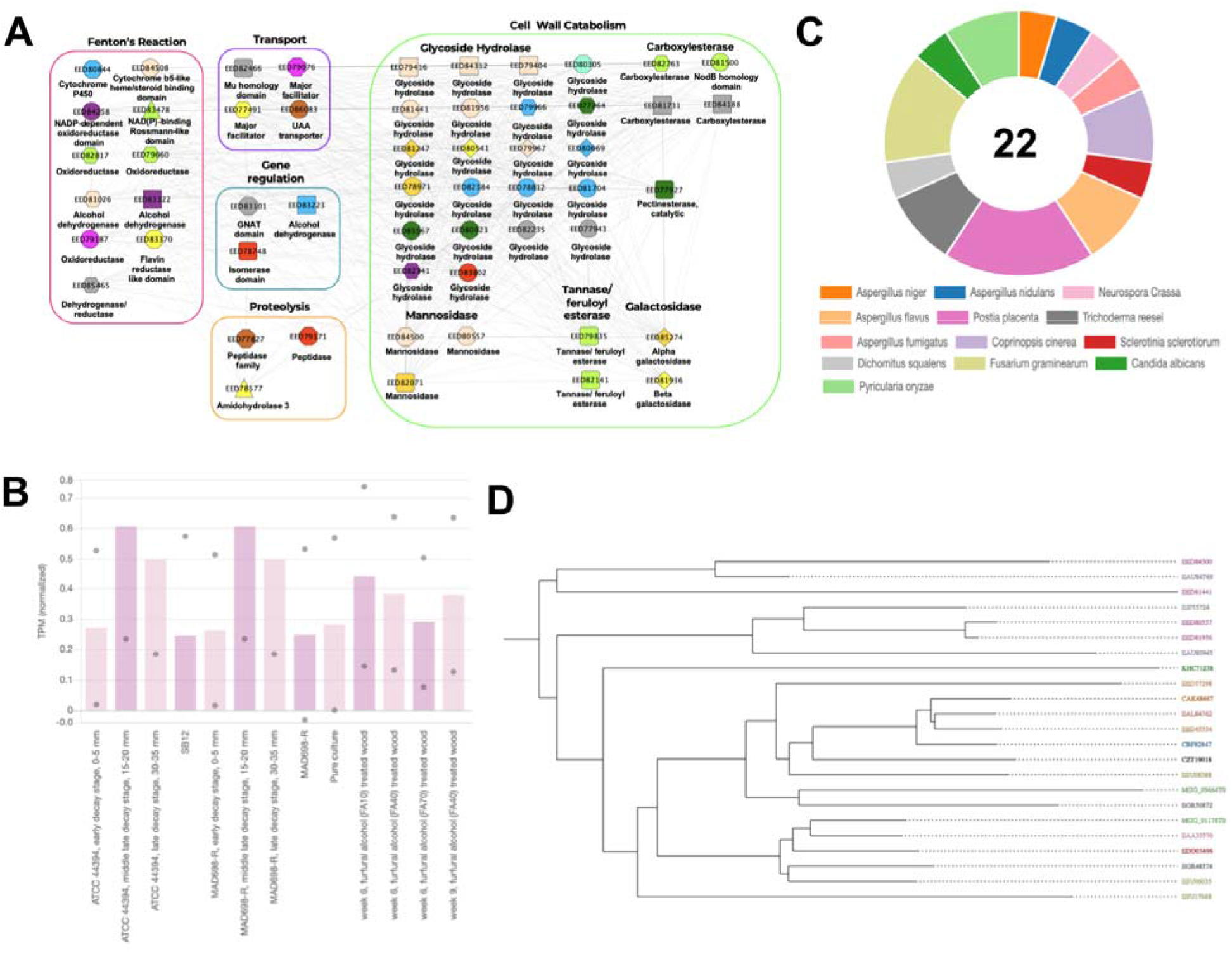
Cluster analysis involved in hydrolase activity in *Postia placenta*. A) Coexpression network of cluster 59 (www.fungi.guru/cluster/view/1745) enriched for hydrolase activity (www.fungi.guru/go/view/10627). Nodes represent genes, while edges connect co-expressed genes. Colored shapes indicate which genes contain Pfam domains and orthogroups in common. The network contains five types of genes: cell wall catabolism genes (indicated in a green rectangle), genes encoding for Fenton’s Reaction (pink rectangle), genes involved in transport (purple rectangle), gene-regulating genes (blue rectangle) and proteolysis associated proteins (orange rectangle). For brevity, only discussed nodes are shown. B) Expression profile of the cluster 59 from POSPL shows that the cluster has the highest expression during the middle-late decay stage in ATCC 44394 and MAD698-R strains C) Proportion of genes in the gene family OG_03_0001073, which is highly abundant in the cluster and which contains genes annotated as glycoside hydrolases and mannosidases.D) Phylogenetic tree of hydrolase gene family OG_03_0001073.

