## Supplementary material for "Fungi.guru: comparative genomic and transcriptomic database for the Fungi kingdom": Table S1

**Table S1. Genome versions used to build the database**

| **Fungal species** | **CDS files** |
| --- | --- |
| *Aspergillus flavus* | Aspergillus_flavus.JCVI-afl1-v2.0.cds.all.fa |
| *Aspergillus fumigatus* | Aspergillus_fumigatus.ASM265v1.cds.all.fa |
| *Aspergillus nidulans* | Aspergillus_nidulans.ASM1142v1.cds.all.fa |
| *Aspergillus niger* | Aspergillus_niger.ASM285v2.cds.all.fa |
| *Candida albicans* | Candida_albicans_sc5314_gca_000784635.Cand_albi_SC5314_V4.cds.all.fa |
| *Coprinopsis cinerea* | Coprinopsis_cinerea_okayama7_130_gca_000182895.CC3.cds.all.fa |
| *Cryptococcus neoformans* | Cryptococcus_neoformans_var_grubii_h99_gca_000149245.CNA3.cds.all.fa |
| *Dichomitus squalens* | Dichomitus_squalens_lyad_421_ss1_gca_000275845.Dichomitus_squalens_v1.0.cds.all.fa |
| *Fusarium graminearum* | Fusarium_graminearum_gca_000240135.ASM24013v3.cds.all.fa |
| *Komagataella phaffii* | (Komagataella_phaffii_gs115_gca_001746955.ASM174695v1.cds.all.fa |
| *Neurospora crassa* | Neurospora_crassa.NC12.cds.all.fa |
| *Postia placenta* | Postia_placenta_mad_698_r_gca_000006255.Postia_placenta_V1.0.cds.all.fa |
| *Puccinia striiformis* | Puccinia_striiformis_gca_002920065.ASM292006v1.cds.all.fa |
| *Pyricularia oryzae* | Magnaporthe_oryzae.MG8.cds.all.fa |
| *Ramularia collo-cygni* | Ramularia_collo_cygni_gca_900074925.version_1.cds.all.fa |
| *Saccharomyces cerevisiae* | Saccharomyces_cerevisiae.R64-1-1.cds.all.fa |
| *Schizosaccharomyces pombe* | Sclerotinia_sclerotiorum.ASM14694v1.cds.all.fa |
| *Sclerotinia sclerotiorum* | Sclerotinia_sclerotiorum.ASM14694v1.cds.all.fa |
| *Trichoderma reesei* | Trichoderma_reesei.GCA_000167675.2.cds.all.fa |
| *Yarrow lipolytica* | Yarrowia_lipolytica.GCA_000002525.1.cds.all.fa |
